# Cell Type Development in *Chlamydia trachomatis* Follows a Program Intrinsic to the Reticulate Body

**DOI:** 10.1101/2020.03.13.991687

**Authors:** Travis J Chiarelli, Nicole A Grieshaber, Anders Omsland, Christopher H Remien, Scott S Grieshaber

**Author notes:** Corresponding Author: Scott Grieshaber, Ph.D., Associate Professor, Department of Biological Sciences, University of Idaho, 875 Perimeter Drive MS 3051, Moscow, ID 83844.

## Abstract

The obligate intracellular bacterial pathogen *Chlamydia trachomatis* (*Ctr*) is reliant on an unusual developmental cycle consisting of two cell forms termed the elementary body (EB) and the reticulate body (RB). The EB is infectious and utilizes a type III secretion system and preformed effector proteins during invasion, but does not replicate. The RB replicates in the host cell but is non-infectious. This developmental cycle is central to chlamydial pathogenesis. In this study we developed mathematical models of the chlamydial developmental cycle that account for potential factors influencing the timing of RB to EB cell type switching during infection. Our models predicted that two broad categories of regulatory signals for RB to EB development could be differentiated experimentally; an “intrinsic” cell autonomous program inherent to each RB or an “extrinsic” environmental signal to which RBs respond. To experimentally differentiate between these hypotheses, we tracked the expression of *Ctr* developmental specific promoters using fluorescent reporters and live cell imaging. These experiments indicated that EB production was not influenced by increased MOI or by superinfection, suggesting the cycle follows an intrinsic program that is not influenced by environmental factors. Additionally, live cell imaging of these promoter constructs revealed that EB development is a multistep process linked to RB growth rate and cell division. The formation of EBs followed a cell type gene expression progression with the promoters for *euo* and *ihtA* active in RBs, while the promoter for *hctA* was active in early EBs/intermediate cells and finally the promoters for the true late genes, *hctB, scc2*, and *tarp* active in the maturing EB.

**Importance:** *Chlamydia trachomatis* is an obligate intracellular bacteria that can cause trachoma, cervicitis, urethritis, salpingitis, and pelvic inflammatory disease. To establish infection in host cells *Chlamydia* must complete a multi cell type developmental cycle. The developmental cycle consists of two specialized cells; the EB which mediates infection of new cells and the RB which replicates and eventually produces more EB cells to mediate the next round of infection. By developing and testing mathematical models to discriminate between two competing hypotheses for the nature of the signal controlling RB to EB cell type switching. We demonstrate that RB to EB development follows a cell autonomous program that does not respond to environmental cues. Additionally, we show that RB to EB development is a function of cell growth and cell division. This study serves to further our understanding of the chlamydial developmental cycle that is central to the bacterium’s pathogenesis.

## Introduction

Chlamydiae are bacterial pathogens responsible for a wide range of diseases in both animal and human hosts (1). *Chlamydia trachomatis* (*Ctr*), a human pathogen, is comprised of over 15 distinct serovars causing both trachoma, the leading cause of preventable blindness, and sexually acquired infections of the urogenital tract (2). According to the CDC, *Ctr* is the most frequently reported sexually transmitted infection in the United States, costing the American healthcare system nearly $2.4 billion annually (3, 4). These infections are widespread among all age groups and ethnic demographics, infecting ~3% of the human population worldwide (5). In women, untreated genital infections can result in pelvic inflammatory disease, ectopic pregnancy, and infertility (6–8). Every year, there are over 4 million new cases of *C. trachomatis* STI infections in the United States (6, 9) and an estimated 92 million cases worldwide (10).

*Chlamydia-related* disease is entirely dependent on the establishment and maintenance of the unique intracellular niche, the chlamydial inclusion, where the bacteria replicate and carry out a biphasic developmental cycle. This cycle generates two unique developmental cell forms: the elementary body (EB) and the reticulate body (RB). The EB cell type mediates host cell invasion via pathogen mediated endocytosis while the RB cell type is replication competent but cannot initiate host cell infection (11). For *C. trachomatis* serovar L2, the cycle begins when the EB binds to a host cell and initiates uptake through the secretion of effector proteins by the Type III secretion system (12). During entry, the EB is engulfed by the host cell plasma membrane forming a vacuole that is actively modified by *Chlamydia* to block interaction with the host endocytic/lysosomal pathway (13). This vacuole, termed the chlamydial inclusion, continues to mature as the EB cell form transitions to the RB cell form. The time from host cell contact to the formation of the mature inclusion containing replication competent RBs is ~11 hours (14). The formation of infectious EB cells occurs reliably between 18-20 hours post infection (HPI) (15). Regulatory control of the transition between the RB and EB is critical for the chlamydial life cycle as *Chlamydia* must balance replication vs. production of infectious progeny. How *Chlamydia* regulates this process is currently unclear although there have been multiple hypotheses proposed to explain the control of the developmental cycle. Regulatory mechanisms such as RB access to, or competition for, inclusion membrane contact (16), reduction in RB size (14) or responses to changes in nutrient availability (17) have all been proposed to control or influence RB to EB cell switching.

In this study we used mathematical modeling to guide experiments to distinguish between factors that influence RB to EB development. The chlamydial life cycle was modeled using systems of differential equations. Each model was tested under simulated conditions which indicated that environmental vs. intrinsic control of EB development could be distinguished experimentally. In order to test the model predictions, a live cell imaging system in combination with promoter reporter constructs was developed to follow the developmental cycle in real time. We show here that neither the limiting membrane hypothesis nor environmental nutrient limiting mechanisms are consistent with our experimental results, and that EB development likely follows a cell autonomous program. Additionally, we show that this intrinsic program is dependent on RB growth and cell division.

## Results

### Modeling chlamydial development

We developed two mathematical models to represent potential driving forces in promoting EB development. The models are systems of ordinary differential equations (ODE) that track RBs, intermediate bodies, and EBs over time (Fig. 1, S1). In these models, development of the EB is controlled by an inhibitory “signal” that is either intrinsic to each bacterium or environmental, i.e., shared between the bacteria (Fig. 1A and B). We did not specify the nature of the signal beyond an inhibitory effect on EB production at high concentrations and it’s consumption by RBs. In each case the signal is consumed by the bacteria over time, and, once depleted, RB to EB conversion commences. The models differ in whether all the RBs in the inclusion compete for one pool of this signal or whether each RB contains an independent internal pool of the inhibitory signal. The output of both models mimic the general kinetics of the chlamydial developmental cycle. Both models produced identical outputs when a multiplicity of infection (MOI) of 1 was simulated (Fig. 1C). When a change in the replication rate of *Chlamydia* was simulated, the two models again responded similarly, showing that an increased replication rate leads to earlier EB production while a decreased replication rate resulted in delayed EB production (Fig. 1D). However, the models produced dramatic kinetic differences with a simulated increase in MOI or time-delayed superinfection. Both simulated conditions caused EB formation to occur sooner in the environmental signal model but had no effect on the intrinsic signal model (Fig. 1E). These data suggest it is possible to experimentally differentiate between whether an environmental signal or an intrinsic program triggers EB development.

**Fig. 1.**
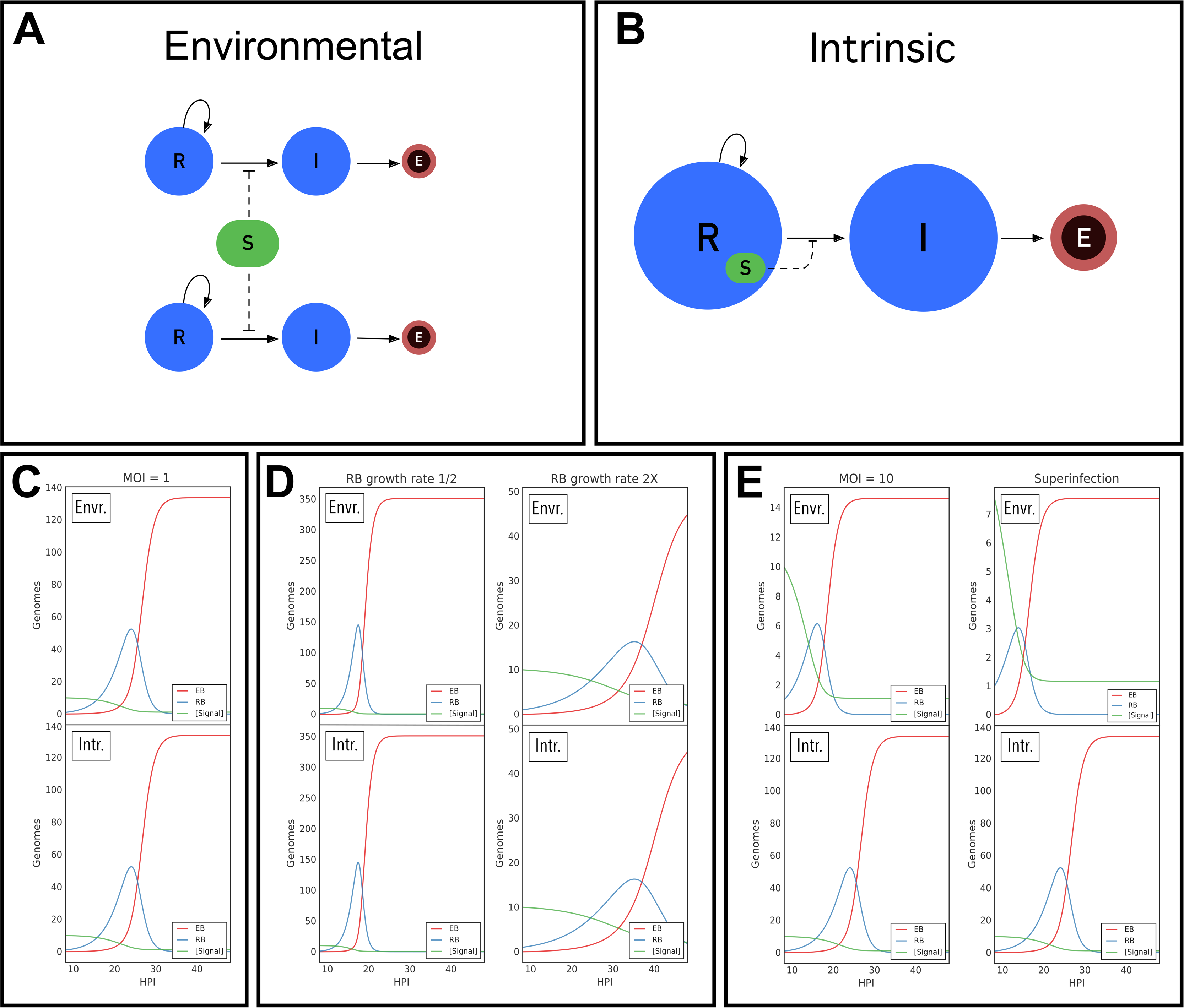
Schematic and simulations of extrinsic and intrinsic models. Both models assume that the mechanism of RB/EB conversion is in response to signal concentration. High signal concentration prevents RB/EB conversion and RB replication continues. As RBs replicate, the signal is consumed and, once the signal is depleted passed a given threshold, RBs convert to IBs (I: blue) which then convert to EBs (E: red). **A.** Schematic of the environmental factor model. The RBs compete for a single pool of signal (S). **B.** Schematic of the intrinsic model. Each RB contains its own signal, eliminating competition between RBs. **C.** Simulations of the two models (extrinsic and intrinsic) using a multiplicity of infection (MOI) of 1 and a RB generation time of 2.27 hours produced results that mimic the kinetics of the chlamydial cycle and were indistinguishable from each other. **D.** Simulations of RB doubling times of 1.13 (½ the measured RB doubling time) hours and 4.54 hours (2x the measured RB doubling time) resulted in a reduced time to EB production and an increased time to EB production, respectively. However, both models produced the same outcome. **E.** Simulations using an MOI of 10 predicted EB conversion to occur more rapidly in the environmental factor model but to remain unchanged in the intrinsic model. Similarly, simulations of the models using a time delayed superinfection resulted in RB to EB conversion occurring more rapidly in the environmental factor model but remaining unchanged in the intrinsic model.

### Development of a live cell reporter system to follow the chlamydial developmental cycle

In order to experimentally differentiate between these possible mechanisms, we developed a live cell imaging system using promoter constructs to follow the chlamydial developmental cycle. The reporter constructs were designed using the promoters of chlamydial genes that are differentially regulated between the RB and EB forms (18). To generate an RB reporter, the promoter of the sRNA IhtA which is expressed early upon infection and negatively regulates the EB specific gene *hctA*, was used to drive EGFP expression (19). To generate an EB reporter, the promoter and first 30nts of the late gene *hctA* was used to drive the expression of the GFP variant Clover. HctA is a small histone like protein that is involved in the condensation of the chlamydial genome to form the compact nucleoid characteristic of the EB (20). The upstream promoter region as well as the first 10 codons of the ORF of HctA were used to construct this reporter as regulation of HctA involves both the promoter and the IhtA binding site contained in the beginning of the ORF (21). Each reporter was transformed into *Ctr*, generating the strains *Ctr ihtA*prom-EGFP and *Ctr hctA*prom-Clover. The chlamydial transformants were used to track the developmental cycle of each strain using live cell time lapse microscopy and particle tracking to quantify the fluorescent expression of individual inclusions over time (22). In order to verify that the fluorescent reporters accurately reflected the developmental cycle, total chlamydial growth was determined by measuring genomic copy by qPCR and EB production by a replating assay to quantify inclusion forming units (IFU). GFP expression from the *ihtA* promoter was first detected at ~10 HPI and started to level off at ~28 HPI (Fig. 2A). The initial expression from the *ihtA* promoter was in good agreement with the initiation of RB cell division as demonstrated by genome copy numbers (Fig. 2A). The initiation of RB replication signals the end of the EB to RB transition after cell entry. Imaging of the *hctA* promoter reporter revealed that the Clover signal could first be detected at ~18 HPI and increased linearly until ~38 HPI (Fig. 2B). Again this was in good agreement with the production of infectious progeny as EBs were first detected at ~20 HPI (Fig. 2B). We measured >50 individual inclusions per strain and found very little inclusion to inclusion variability in the timing of initiation of expression (Fig. 2C and D). This uniformity in developmental timing can be appreciated in a live cell time lapse movie of *Ctr hctA*prom-Clover (Fig. M1). The close agreement between classic methods for following the chlamydial developmental cycle (IFU and genomic copy number) and the single inclusion based fluorescent reporter system described here demonstrates that this system accurately reflects the developmental cycle.

**Fig. 2.**
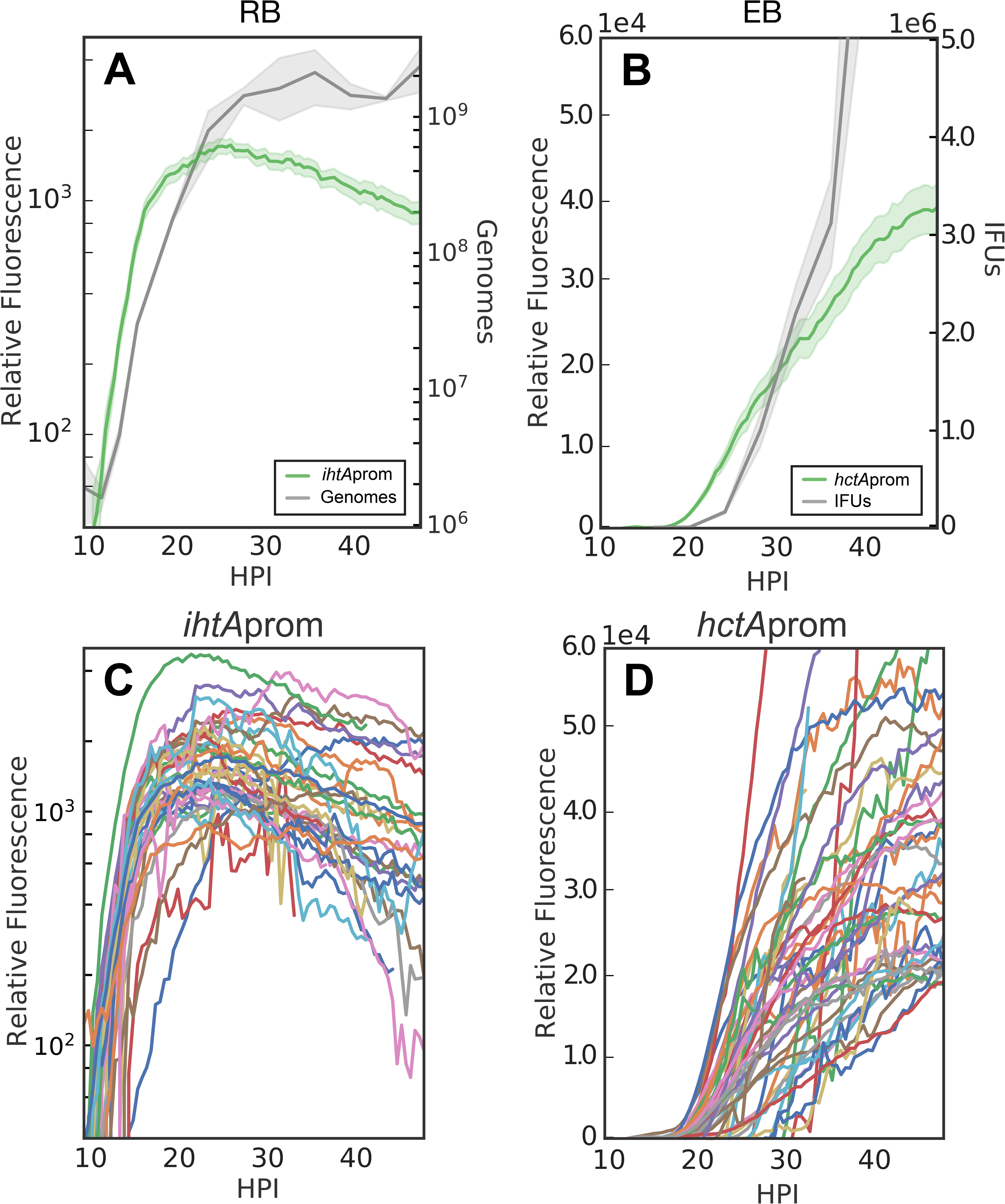
Live-cell fluorescent imaging of chlamydial development. Cell-type specific fluorescent reporters were created to track chlamydial development in real-time. Infections with purified *Ctr-*L2-prom EBs were synchronized and fluorescence microscopy and GE/IFU experiments were run simultaneously. **A-B.** The average of *ihtA*prom-EGFP (RB) and *hctA*prom-Clover (EB) expression intensities from >50 individual inclusions monitored via automated live-cell fluorescence microscopy throughout the developmental cycle compared to genome copy number and IFU, respectively. **C-D.** The fluorescence intensities of >50 individual inclusion tracked via live-cell microscopy throughout the developmental cycle indicated that initiation of fluorescence was fairly uniform. GE and IFU cloud represents 95% ci, fluorescent intensity cloud represents SEM.

### Chlamydial development is growth rate dependent

Both models predicted that changes in growth rate would be reflected in EB production times and rates (Fig. 1D). There is generally a linear relationship between temperature and the square root of growth rate in bacteria (23). Therefore, to validate the predictions of our two models, we followed the live cell reporters at three temperatures; 35°C, 37°C (control), and 40°C. As expected, at the lower temperature of 35°C the EB to RB lag time increased dramatically and *ihtA*prom-EGFP expression increased more slowly than the 37°C control (Fig. 3A). The lower replication rate at 35°C was also reflected in the qPCR genome counts (Fig. 3B). Conversely, the lag time to fluorescence detection was reduced and increased faster than the control when grown at 40°C (Fig. 3A). As predicted by our models, time to EB production was also shifted by changes in growth rate as *hctA*prom-Clover expression began earlier at 40°C and was delayed at 35°C (Fig. 3C). These results were verified by measuring the production of infectious progeny (Fig. 3D). Taken together, these data provide strong evidence that the cycle is growth rate dependent and that our experimental system accurately detected changes in chlamydial development.

**Fig. 3.**
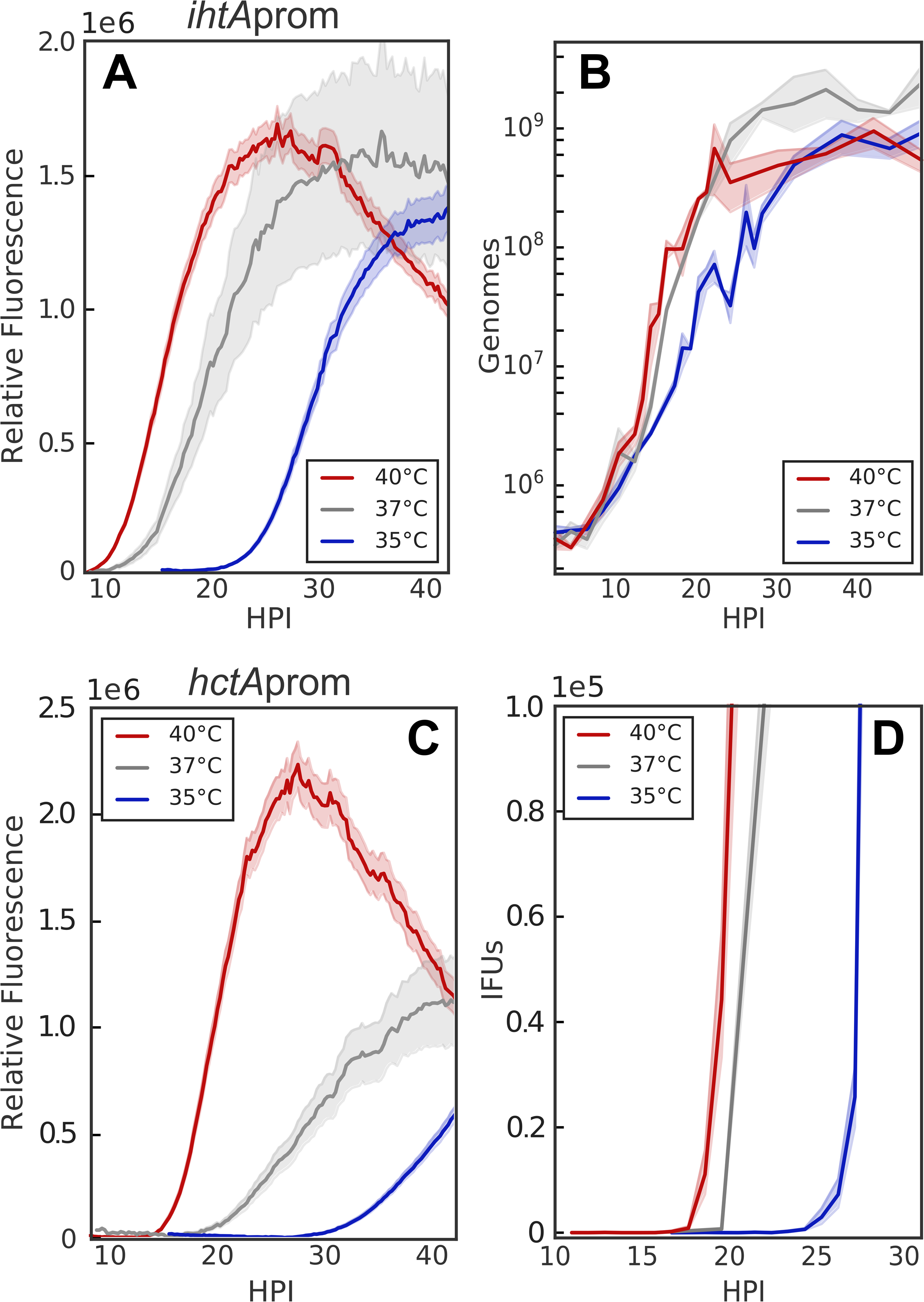
RB replication and EB conversion are growth rate dependent. The ability of the promoter reporter system to monitor differences in RB replication and EB conversion was tested by altering the growth temperature (35°C: blue, 37°C: grey, and 40°C: red). **A-B.** The average of *ihtA*prom-EGFP (RB) and *hctA*prom-Clover (EB) expression intensities from >50 individual inclusions monitored via live-cell fluorescence microscopy from 9-48 HPI. **C.** GEs were measured hourly between 9 and 48 HPI by qPCR. **D.** EB conversion (IFU) was quantified via reinfection assay. Each set of experiments was run in triplicate. GE and IFU cloud represents 95% ci, fluorescent intensity cloud represents SEM.

### EB development is controlled by intrinsic factors and not environmental factors

The two mathematical models differ principally in the source of the EB development signal: internal vs. environmental. The models produced divergent outcomes under conditions where bacteria are competing for a host cell or intra-inclusion signal vs. a signal internal to each RB. Simulations predicted that the time to EB production would be measurably affected by increasing the MOI if the signal was environmental (competitively consumed) but would be unchanged if the signal was intrinsic (internal to each RB) (Fig. 1E). To measure this we used live cell imaging to assess the effect of MOI on EB production. To more accurately assay EB development, two additional EB gene reporters were constructed. The promoters and first 30nts of *hctB*, and *scc2* were inserted upstream of Clover. Like HctA, HctB is a small histone-like protein that is involved in EB nucleoid formation (24) while Scc2 is a chaperone for Type III secretion effector proteins (25). Our published RNA-seq data showed that the transcripts for *hctB* and *scc2* were expressed late, corresponding to the timing of EB production (18). Cells were infected with MOIs ranging from 1 to 32 infectious EBs per cell and imaged every 30 minutes for 40 hours. The MOI was calculated by infecting with a two-fold dilution series and back calculating from an MOI of 1. The fluorescent signals were normalized by MOI. Expression initiation of the RB reporter, *ihtA*prom, did not vary as a function of MOI (Fig. 4A). Expression initiation of the EB promoter reporters, *hctA*prom, *hctBprom* and *scc2*prom, also did not vary as a function of MOI (Fig. 4B-D). The lack of MOI response for the expression of EB genes corresponds closely with EB production as measured by a reinfection assay (Fig. 4E). We also noticed a dramatic difference in the timing of expression between the late genes; *hctA*prom-Clover expression was initiated at ~18 hour post infection while *hctB*prom-Clover and *scc2*prom-Clover expression was initiated ~3 hours later at ~ 21 hour post infection (HPI).

**Fig. 4.**
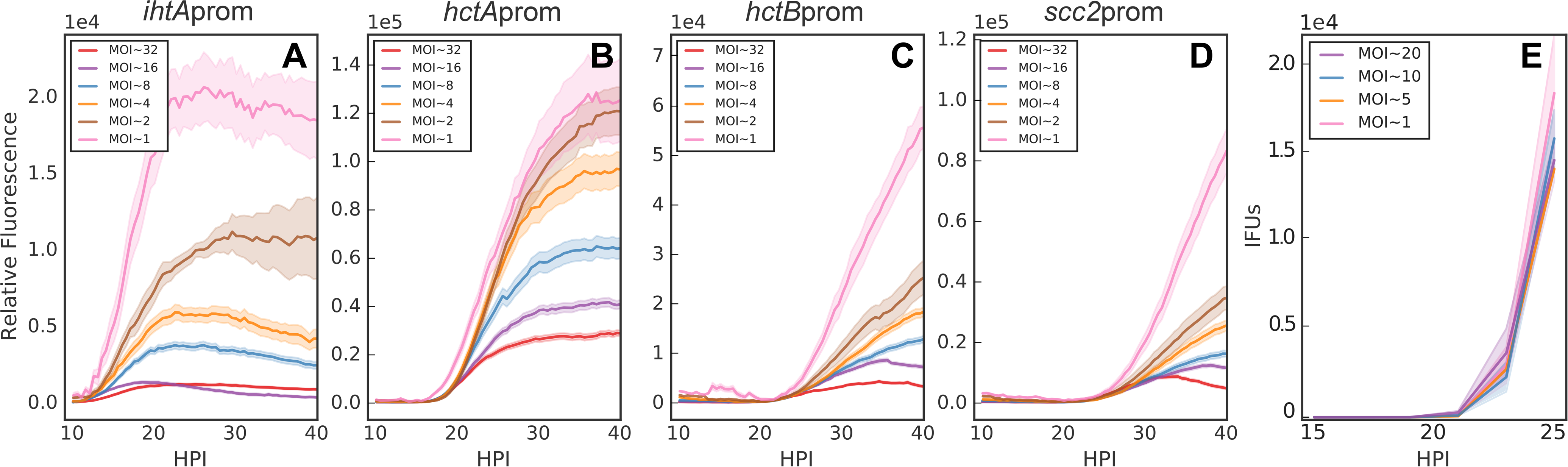
MOI does not affect RB to EB conversion. Host cells were infected with purified *Ctr-*L2-prom EBs at an MOI of 1-32. **A-D:** The average of *ihtA*prom-EGFP (RB) and *hctA*prom-Clover, *hctB*prom-Clover, and *scc2*prom-Clover (EB) expression intensities from >50 individual inclusions monitored via automated live-cell fluorescence microscopy throughout the developmental cycle. Fluorescent intensities were normalized by respective MOI. **E.** EB development (IFU) was measured at MOIs from 1-20 and was quantified via a reinfection assay. EBs were harvested at two-hour intervals from 15-25 HPI. IFU data was normalized by respective MOI. Each set of experiments was run in triplicate. IFU cloud represents 95% ci, fluorescent intensity cloud represents SEM.

Our models predicted that both MOI and superinfection conditions would differentiate between cell autonomous development and environmentally influenced development (Fig. 1E). The MOI data suggested that RB to EB developmental switching is not influenced by the host intra-cellular or intra-inclusion environment but rather, is triggered by an intrinsic signal. To further differentiate between these possibilities, we measured RB and EB gene expression under superinfection conditions. The chlamydial inclusion is derived from the plasma membrane and interaction with the endocytic membrane system is actively blocked by *Chlamydia* (13). When multiple EBs infect a cell they each create individual inclusions that traffic to the microtubule-organizing center (MTOC) of the cell (26). This trafficking, along with the expression of the fusogenic protein IncA at ~12 HPI, culminates in homotypic inclusion fusion resulting in a single chlamydial inclusion per cell (27, 28). Our extrinsic model predicted that the developmental cycle of *Chlamydia* under superinfection conditions would be dramatically altered (decreased time to EB production) as a function of the developmental stage of the initial infection. To test this, cells were infected with unlabeled *Ctr* L2 for either 6, 12 or 18 hrs prior to a second infection with *Ctr* L2 containing the reporter plasmids, and imaged starting at 9 hrs post secondary infection. Fluorescent signals were measured for inclusions that were verified to be superinfected by imaging both DIC and fluorescence, i.e. inclusions containing both labeled and unlabeled *Chlamydia* (Fig. 5A). Superinfection at any time post initial infection had no effect on the timing of the expression of either *ihtA*prom-EGFP or *hctA*prom-Clover (Fig. 5B and C). Although the time of gene expression was unchanged, there did appear to be an effect on carrying capacity that was time-dependent, suggesting an upper limit in total EB production per cell (Fig. 5C). The lack of effect on late gene expression was verified with two other late promoter reporters, *hctB*prom-Clover and *scc2*prom-Clover, twelve hours post superinfection (Fig. 5D and E). We verified that superinfection had no effect on the production of infectious progeny by performing a replating assay in the presence of spectinomycin (Fig. 5F).

**Fig. 5.**
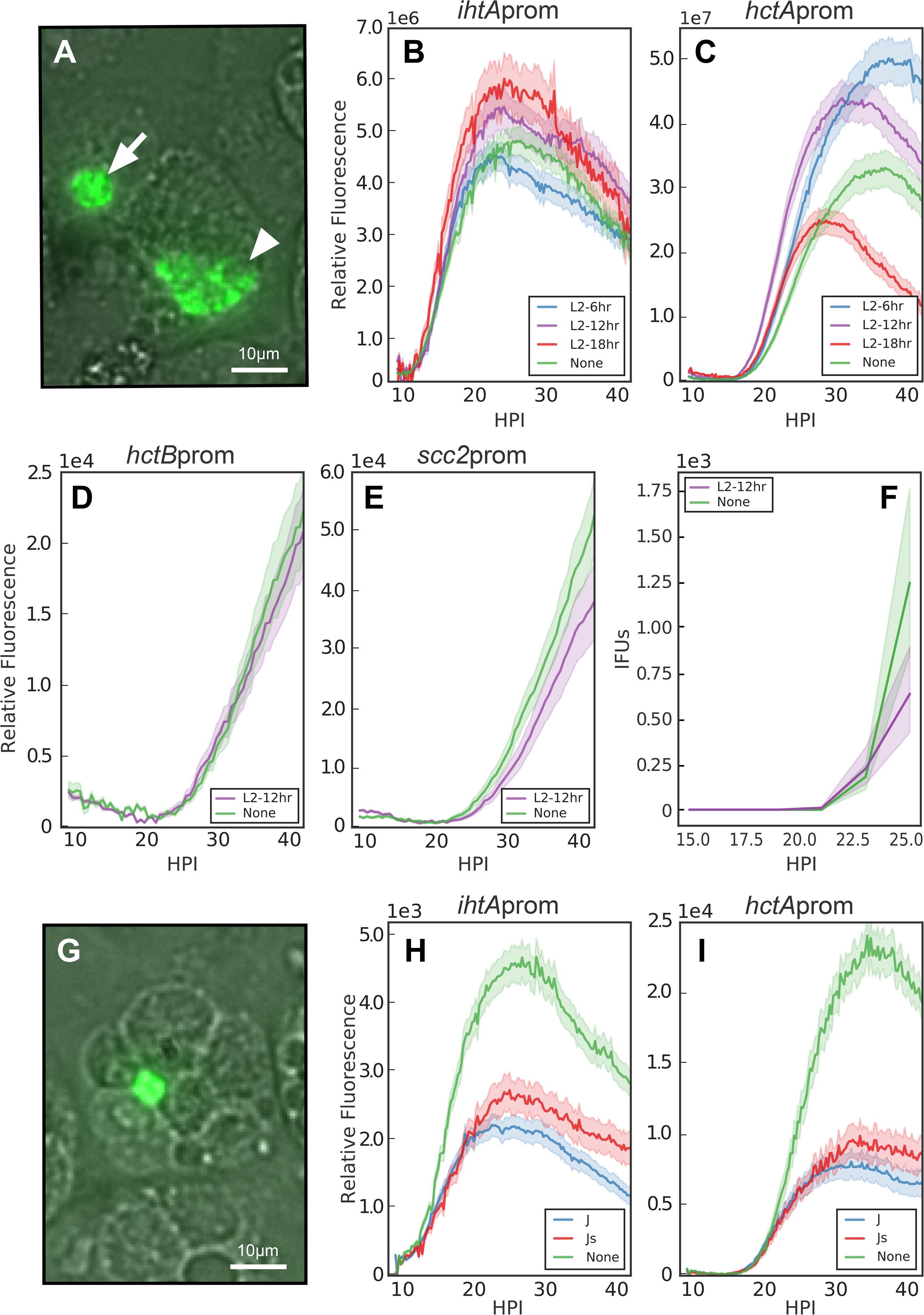
Superinfection does not affect RB to EB conversion. Host cells were initially infected with non-fluorescent *Ctr* EBs followed by secondary infections with *Ctr-*L2-prom EBs at 6, 12, or 18 HPI and the fluorescent output compared to cells that had not been infected with a primary infection (None). Infections were imaged starting at 9 hours post infection with the *Ctr-*L2-prom strains. **A.** Live-cell fluorescence/DIC image of 18 hr L2 superinfection with *Ctr*-L2-*hctA*prom-Clover at 20X magnification (30 hours post *Ctr*-L2-*hctA*prom infection). Fluorescent signals were measured in inclusions containing both GFP expressing *Ctr* (arrowhead) and non fluorescent *Ctr* (arrow). Scale bar = 10μm. **B-C.** The average of *ihtA*prom-EGFP (RB) and *hctA*prom-Clover (EB) expression intensities from >50 individual inclusions monitored via automated live-cell fluorescence microscopy during no superinfection, 6, 12, and 18 hour *Ctr*-L2 superinfections. **D-E.** The average fluorescent intensities of >50 individual inclusions using *Ctr*-L2-*hctB*prom-Clover or *Ctr-*L2-*scc2*prom-Clover measured with no superinfection (None) or 12 hour *Ctr*-L2 superinfection. **F.** EBs were harvested at two-hour intervals from 15-25 hours post *Ctr-*L2-prom infection and quantified by reinfection assay. **G.** Live-cell fluorescence/DIC image of cells infected with Js and superinfected with *Ctr*-L2-*hctA*prom-Clover. Image was taken 30 hours post *Ctr*-L2-*hctA*prom-Clover infection at 20X magnification. Fluorescent signals were measured from inclusions in cells that contained both fluorescent *Ctr* (arrowhead) and unfused non fluorescent *Ctr* (arrow). Scale bar = 10μm. **H-I.** The average of >50 individual inclusions containing *ihtA*prom-EGFP (RB) and *hctA*prom-Clover (EB) fluorescent intensity measured with no superinfection (None), *Ctr*-J, or *Ctr*-Js superinfections. Each set of experiments was repeated three times. IFU cloud represents 95% ci, fluorescent intensity cloud represents SEM.

To further examine any effect of the intra-inclusion environment versus the host intra-cellular environment, we took advantage of a *Chlamydia* mutant that does not express IncA and is therefore defective in homotypic inclusion fusion (27). Cells were pre-infected with an isogenic mutant pair, either *Ctr* J (incA positive and fusogenic (29)) or *Ctr* Js (incA negative and non fusogenic (29)) for 18 hours and were then superinfected with *Ctr* L2 containing the reporter plasmids (*Ctr* L2 prom) and imaged starting at 10 hrs post superinfection (Fig. 5G). Again, there was no apparent change in growth rate for either infection alone (no superinfection), superinfection with inclusion fusion or superinfection without fusion (Fig. 5H and I). Interestingly, total signal for the late genes was reduced for both *Ctr* Js and *Ctr* J superinfections as compared to *Ctr* L2 prom strains alone suggesting that carrying capacity is likely defined by the host intra-cellular environment and not the intra-inclusion environment (Fig. 5H and I). Taken together these data suggest that the timing of RB to EB development is an intrinsic pre-programmed property of *Chlamydia* and does not respond to environmental signals.

### Cell division is required for EB development

Time to EB development responded to RB growth rate suggesting that cell division may be critical for development (Fig. 3). To test the role of cell division in EB development, RB replication was halted by treating infected cells with penicillin (Pen). *C. trachomatis* does not use peptidoglycan as a structural sacculus and *Ctr* does not contain a peptidoglycan cell wall. Instead, peptidoglycan aids cell septation by forming a ring at the cleavage furrow (30). Therefore, Pen treatment blocks cell septation but not cell growth. Cells infected with *Ctr ihtA*prom-EGFP or *Ctr hctA*prom-Clover were treated with Pen at 14 HPI and imaged for a further 34 hrs. The fluorescent signal for individual inclusions was measured using single inclusion tracking. The *ihtA*prom signal after Pen treatment continued to increase as did the size of the RB cells (Fig. 6A and Fig. 8A-B). This increased signal intensity initially tracked that of untreated cells but the fluorescence intensity in Pen treated inclusions increased more rapidly as compared to untreated (Fig. 6A). *IhtAprom* expression in the presence of Pen also matched the increase in genome copies which, as previously reported (31), was also Pen insensitive (Fig. 6B). Unlike *ihtA*prom-EGFP expression, the *hctA*prom-Clover signal was dramatically affected by Pen treatment. *hctA*prom expression was initially repressed by Pen treatment at 14 HPI, however expression was initiated ~9 hrs after treatment (Fig. 6C and D). We explored this expression behavior further using three other late gene promoter reporters, *hctB*prom-Clover, *scc2*prom-Clover and *tarp*prom-Clover (Fig. 6C-D and S2). The expression pattern driven by *hctB*prom, *scc2*prom and *tarp*prom were dramatically different from *hctA*prom as none showed Clover expression in the Pen treated samples (Fig. 6D and S2). The lack of *hctB*prom, *scc2*prom and *tarp*prom gene expression corresponded with the lack of production of infectious progeny during Pen treatment suggesting that these genes can be considered true “EB” genes (Fig. 6E).

**Fig. 6.**
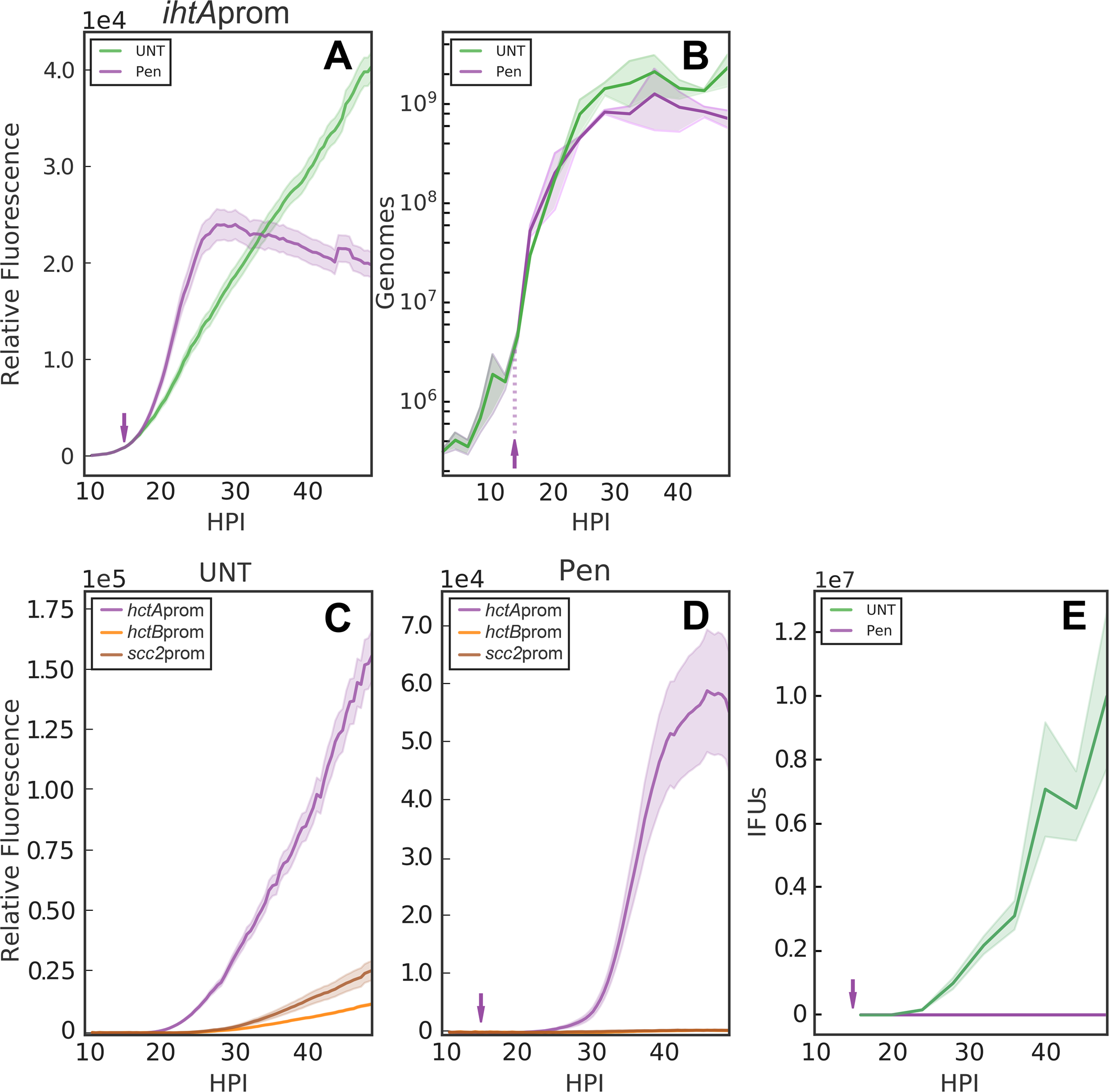
Inhibition of cell division inhibits EB conversion. Host cells were infected with purified *Ctr-*L2-prom EBs followed by addition of penicillin-G at 14HPI (arrow) or vehicle only. **A.** The average of *ihtA*prom-EGFP (RB) expression intensities from >50 individual inclusions monitored via automated live-cell fluorescence microscopy in the absence (UNT) or presence of penicillin (Pen). **B.** Genome copy numbers measured using qPCR. **C-D.** The average of *hctA*prom-Clover*, hctB*prom-Clover and *scc2*prom-Clover (EB) fluorescence intensities from >50 individual inclusions monitored via automated live-cell fluorescence microscopy in the absence (UNT) or the presence of penicillin (Pen). **E.** EBs were harvested at four-hour intervals for each treatment and quantified by reinfection assay. Each set of experiments was repeated four times. GE and IFU cloud represent 95% ci, fluorescent intensity cloud represents SEM.

To further investigate the role of cell division in EB development, we tested the effects of a second antibiotic that targets peptidoglycan synthesis, D-cycloserine (DCS). DCS is a cyclic analogue of D-alanine and acts against two enzymes important in peptidoglycan synthesis: alanine racemase (Alr) and D-alanine:D-alanine ligase (Ddl) (32). GFP expression in single inclusions was measured over time after DCS treatment at 14 HPI. An additional early gene promoter reporter, *euo*prom-Clover, was included to explore RB expression behavior further. EUO (early upstream ORF) is a transcriptional repressor that selectively regulates promoters of *Ctr* late genes (33). Both early gene promoter strains treated with DCS expressed fluorescence with similar kinetics to Pen treated and untreated samples (Fig. 7A-C). The expression kinetics of the late gene reporters after DCS treatment also mimicked Pen treatment. DCS treated inclusions never expressed Clover from *hctB*prom or *scc2*prom reporters but did express from the *hctA*prom reporter with a similar ~9 hour delay (Fig. 7 D-F). Although the kinetics were similar to Pen treatment among all reporters, the aberrant RBs did not grow as large as those treated with Pen (Fig. 8B and C).

**Fig. 7.**
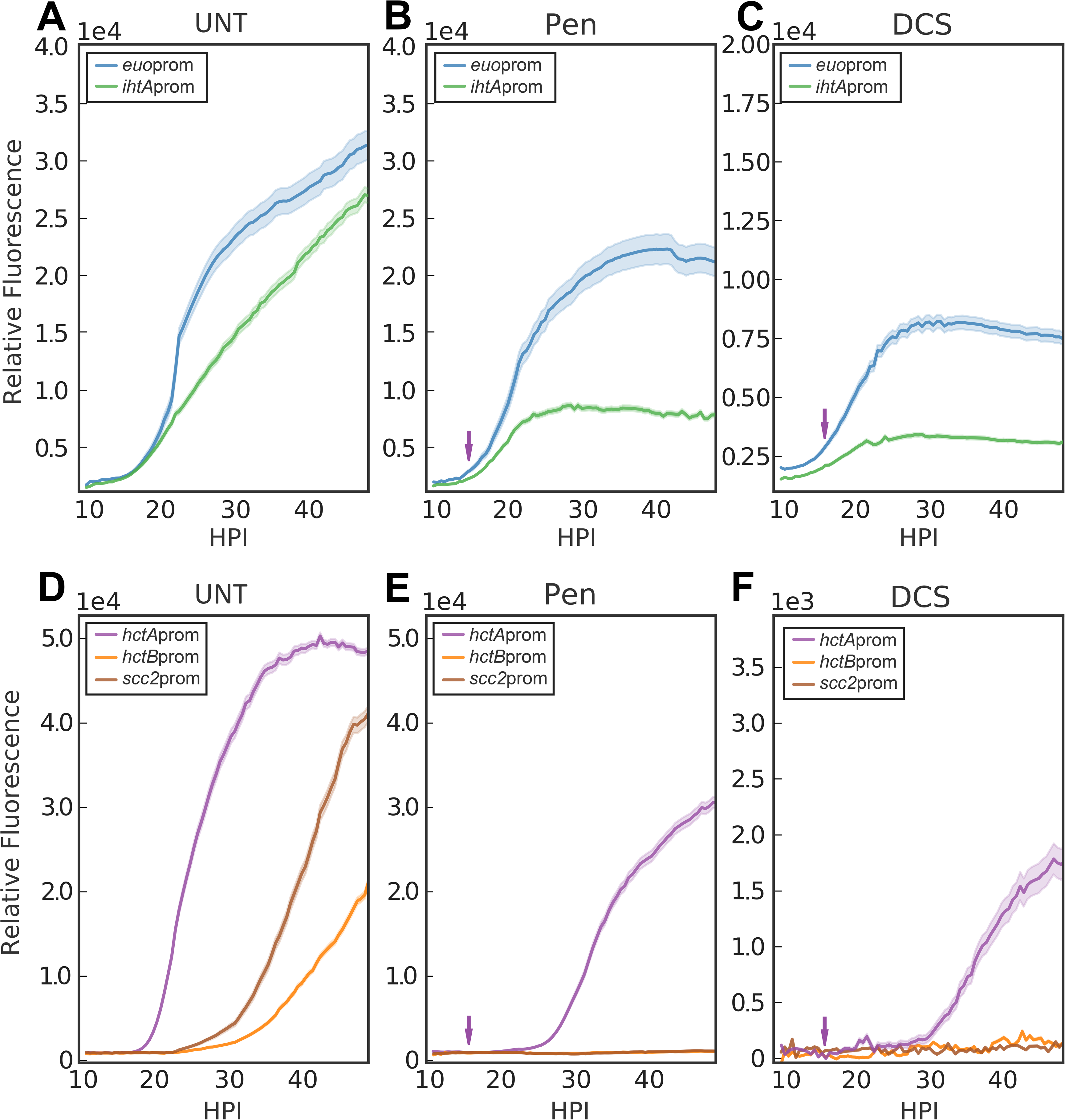
Effects of an alternative developmental inhibitor on RB/EB conversion. Host cells were infected with *Ctr*-L2-prom EBs followed by treatment with penicillin-G, D-cycloserine, or vehicle only at 14HPI. **A-C.** The average of RB (*ihtA*prom-EGFP, *euo*prom-Clover) expression intensities from >50 individual inclusions monitored via automated live-cell fluorescence microscopy in vehicle only (UNT), penicillin (PEN), or D-cycloserine (DCS) treated cells, respectively. **D-F.** The average of *hctA*prom-Clover, *hctB*prom-Clover and *scc2*prom-Clover (EB) fluorescent intensities from >50 individual inclusions in UNT, PEN, or DCS treated cells, respectively. Each set of experiments was repeated four times. Cloud represents SEM.

**Fig. 8.**
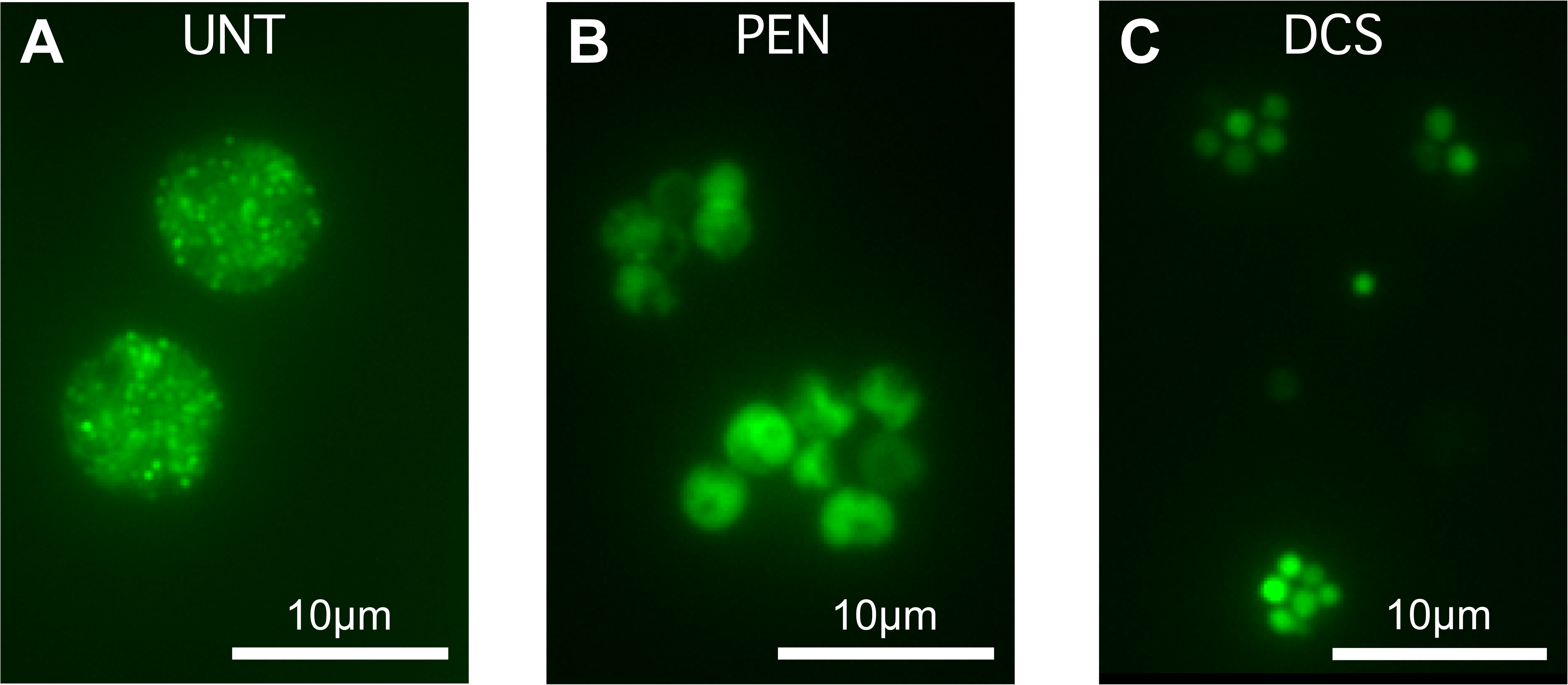
D-cycloserine and penicillin-G induces aberrant cell forms. Host cells were infected with Ctr-L2-*hctA*prom-Clover EBs followed by treatment with PEN or DCS at 14HPI. Live-cell fluorescence images were aquired at 40 HPI. A) Untreated, vehicle only **B**) Penicillin treated (PEN) and **C**) D-cycloserine treated (DCS). Magnification, 40X. Scale bar = 10μm.

Treatment with penicillin has been reported to induce a metabolic state termed persistence as the aberrant RBs continue to metabolize but do not produce infectious progeny (34, 35). Pen, other antibiotic treatments, and nutrient limitation are all reported to induce a persistent growth state in *Chlamydia* (36). Therefore we explored the effect of interferon-gamma (IFN-γ) induced persistence on cell type gene expression. While Pen and DCS induce persistence through their effects on peptidoglycan synthesis, IFN-γ causes a persistent state by starving *Chlamydia* of tryptophan (37). HeLa cells were used as opposed to Cos7 cells as they respond well to hIFN-γ. Cells were treated with IFN-γ 24 hrs prior to infection with *Ctr ihtAprom-EGFP* or *Ctr* HctAprom-Clover strains. Imaging of these constructs showed that no signal was produced from either promoter construct after IFN-γ treatment suggesting a dramatic reduction in overall metabolic activity (Fig 9).

**Fig. 9.**
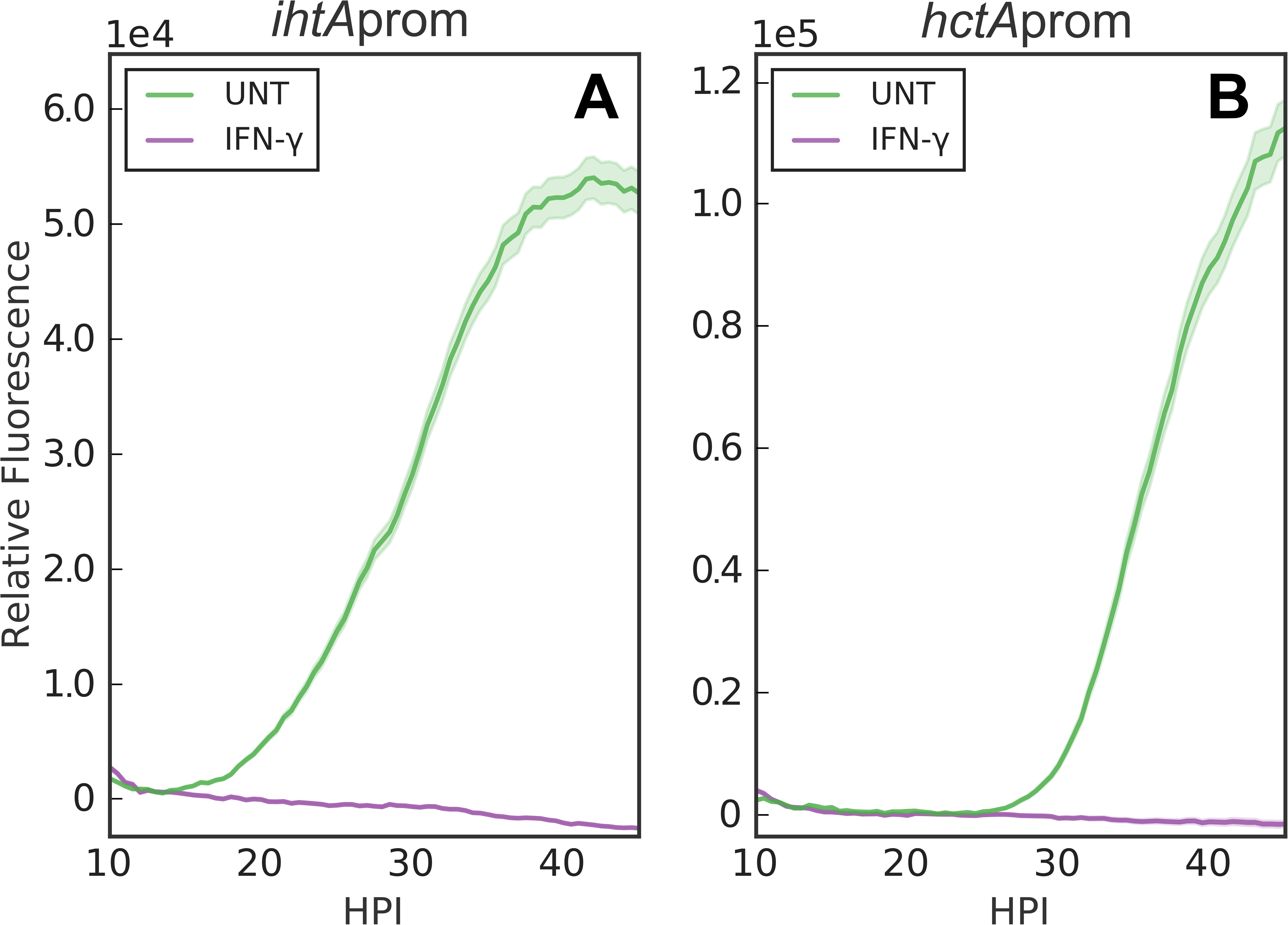
Effect of interferon-gamma on RB/EB conversion. HeLa cells were treated with interferon-gamma (IFN-γ) 24 hrs prior to infection with *Ctr-*L2-prom EBs. **A**. The average of *ihtA*prom-EGFP (RB) expression intensities from >50 individual inclusions monitored via automated live-cell fluorescence microscopy in the absence (UNT) and presence of IFN-γ. **B.** Plots represent the average *hctA*prom-Clover (EB) fluorescent expression from >50 individual inclusions. Each set of experiments was repeated four times, cloud represents SEM.

The Pen and DCS data support a role for cell division in chlamydial development. To further explore this observation, cells were treated with Pen every two hours starting at 16 HPI. To visualize both RBs and EBs in the same inclusion during the developmental cycle, two dual promoter constructs were developed, *hctA*prom-mKate2/*ihtA*prom-mNeonGreen and *hctB*prom-mKate2/*euo*prom-Clover. Cells were infected with the dual promoter strains and imaged every 30 minutes after addition of Pen at the indicated times (Fig. 10). The responses of the promoters to Pen treatment were strikingly different. The *euo*prom signal increased in comparison to untreated infections almost immediately after Pen was added, regardless of timing of treatment (Fig. 10A). The *hctB*prom signal was completely inhibited but only after an ~10 hour delay, again regardless of when Pen was added (Fig. 10B). Conversely, *hctA*prom signal was inhibited very quickly after Pen treatment but expression resumed after an ~ 9 hour delay (Fig. 10C). Pen treatment blocks chlamydial cell septation resulting in the formation of large aberrant cells. Confocal images of Pen treated cells indicated that *ihtA*prom-mNeonGreen and *euo*prom-Clover expression was evident only in the large aberrant cells (Fig. 11). However, there was a striking difference in cell type expression location between the late promoters *hctA*prom and *hctB*prom. Like *ihtAprom* and *euo*prom, *hctA*prom-mKate2 expression was localized to the large aberrant cells. In contrast, *hctB*prom-mKate2 expression was restricted to non-aberrant small cells that resembled EBs (Fig. 11).

**Fig. 10.**
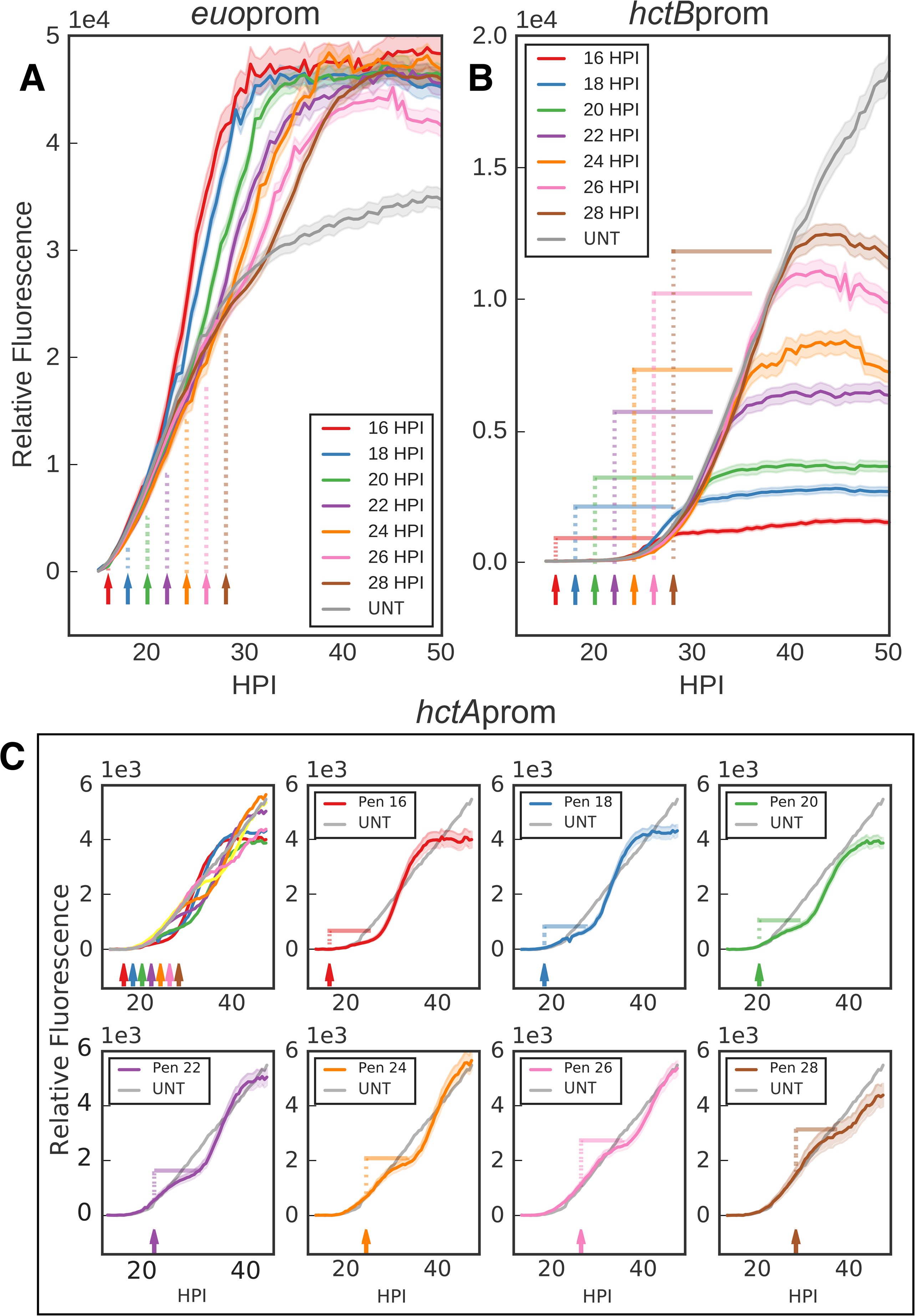
Inhibiting cell-division inhibits further EB conversion. Host cells were infected with *Ctr-L2-prom* EBs followed by treatment with Pen at two-hour intervals starting at 16 HPI or without treatment (UNT). Arrows and vertical dotted lines indicate the addition of Pen. **A.** The average of *euo*prom-Clover (RB) expression intensities from >50 individual inclusions monitored via automated live-cell fluorescence microscopy for each penicillin treatment (time series starting at 16HPI) and untreated (UNT). **B.** The average of *hctB*prom-mKate2 (EB) fluorescence from >50 individual inclusions. Horizontal solid lines indicate time to maximum expression. **C.** The average of *hctA*prom-mKate2 (EB) fluorescence from >50 individual inclusions. *HctA*prom-mKate2 graphs are separated for clarity. Horizontal solid lines indicate time to reinitiation of expression. Each set of experiments was repeated three times. Cloud represents SEM.

**Fig. 11.**
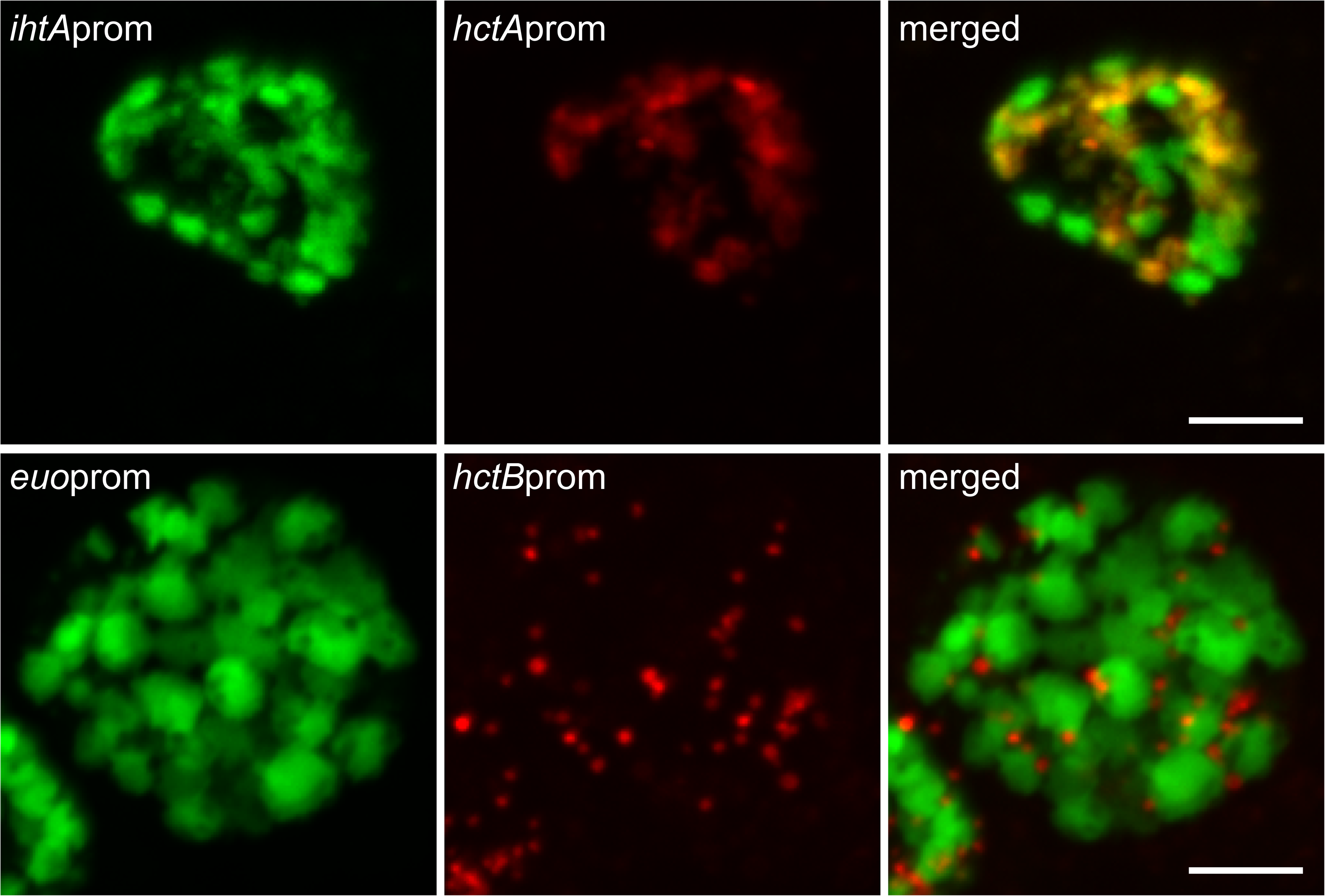
Confocal fluorescence microscopy of cell-type promoter expression upon inhibition of cell division. Host cells were infected with *Ctr-*L2-*hctA*prom-mKate2/*ihtA*prom-mNeongreen (red and green respectively) or *Ctr*-L2-*hctB*prom-mKate2/*euo*prom-Clover (red and green respectively) followed by treatment with Pen at 20 HPI. Samples were fixed at 24HPI. Fixed samples were imaged by confocal microscopy and maximum intensity projections are shown. Scale bars = 5μm.

## Discussion

Infection of vertebrate hosts by *Chlamydia* is dependent on the transition between two specific cell types, the RB and EB, that each have specialized functions. The RB undergoes cell division although it is not yet clear whether this division occurs by binary fission (14) or polarized division (38). The EB form is responsible for mediating invasion of eukaryotic host cells and does not undergo cell division. The EB does however metabolize nutrients to maintain its infectious phenotype (18). This division of labor presents an interesting dilemma for *Chlamydia* as increasing cell numbers through RB division must be balanced with production of infectious EBs. How *Chlamydia* regulates this balance is currently unknown.

Proposed mechanisms for the control of RB to EB development can be divided into two broad categories; a response to extrinsic environmental cues or an intrinsic developmental program. By developing mathematical models and running simulations of infection conditions, we determined that these two possibilities could be differentiated by generating competition between RBs for environmental signals or nutrients. To explore these models experimentally, we developed a live cell reporter system to follow cell type switching in real time at the single inclusion level. Cell-type specific promoters were used to drive the expression of fluorescent proteins in order to follow RB growth (*ihtA*prom and *euo*prom) and EB development (*hctA*prom, *hctB*prom, *scc2*prom and *tarp*prom). Growth and development curves generated using the live cell reporter constructs were comparable to developmental data generated using qPCR and reinfection assays.

Competition for nutrients by increasing MOI and time delayed superinfections, both of which generate competition for host cell and intra-inclusion signals, did not alter time to EB development. These data strongly suggest that development from RB to EB is independent of the intra-inclusion or host environment, but rather is responsive to one or more intrinsic cell autonomous signals. Although timing of RB to EB development was unchanged by environmental conditions, a reduction in maximum *hctA*prom gene expression in the L2 18hr superinfection and serovar J and Js superinfection experiments was notable, indicating that infected cells have an overall carrying capacity. Interestingly, total *hctA*prom gene expression during *Ctr* J (fusogenic) and *Ctr* Js (non fusogenic) superinfections were similar suggesting that the carrying capacity is controlled at the host cell level.

The use of live cell promoter reporters to interrogate cell type switching dramatically improved the resolution for following development. Reporter expression was measured every 30 minutes at the single inclusion level which led to the identification of two different classes of late promoters. *hctB, scc2* and *tarp* were all expressed ~22 HPI, and are therefore considered a class of true late genes. However, our data suggest that *hctA* should be considered an early late gene as its promoter is induced hours before the other late genes tested and responds differently to inhibition of cell division. This differential timing in expression between HctA and the late genes is corroborated by our published RNA-seq data that demonstrated that the transcript encoding HctA was upregulated at 18 HPI while the transcripts for HctB, Scc2 and Tarp were not detected until 24 HPI (18).

Our data also showed that the developmental program is linked to both growth rate and cell division. *Chlamydia* grown at 35°C replicated slower and EB development was delayed compared to those grown at 37°C. Conversely, *Chlamydia* incubated at 40°C replicated faster and EB development was initiated earlier compared to those grown at 37°C. In addition to growth, cell division was also required to trigger EB development. Penicillin and DCS both target peptidoglycan synthesis at different points in the pathway resulting in a block in cell septation during chlamydial replication (39). Both treatments, when added early in infection (prior to 12 HPI), inhibited EB formation as measured by production of infectious particles and expression of late gene promoter reporters (*hctA*, *hctB*, *scc2*, and *tarp*). However, the effect of these drugs on *hctA*prom-Clover expression differed significantly from the effects seen on *hctB*, *scc2*, and *tarp*. Although *hctA*prom-Clover expression was initially inhibited, expression was eventually initiated in the aberrant forms after an approximate 9 hr delay.

Pen addition at all times tested (two hour intervals from 16h-28h) resulted in an immediate overall increase in *euo*prom-Clover expression and an immediate overall decrease in *hctA*prom-Clover expression in inclusions compared to untreated samples. In contrast, *hctB*prom expression kinetics was similar to untreated controls for approximately 10 h post Pen addition, at which point further expression was inhibited. Additionally, *hctB*prom fluorescence was only evident in small cell forms indicating expression was restricted to EBs while *hctA*prom expression was evident in the RB like aberrant forms suggesting expression in an intermediate cell form. These data suggest that inhibiting cell division blocks RBs from switching off *euo*prom expression and switching on *hctA*prom gene expression. However, if a cell is already committed to EB formation (*hctA*prom positive), EB gene expression continues (Pen insensitive) until the EB is fully mature (maximal *hctB*prom signal), which our data indicates takes about 10 hours.

Treatment of *Chlamydia*-infected cells with antibiotics or nutrient limitation causes a growth phenotype termed persistence (36). Persistence is characterized by aberrant RB forms which are larger in comparison to untreated RBs, do not undergo cell division, and do not produce infectious progeny (36). Although all these treatments cause aberrant RBs, the phenotypes vary (37, 40). Pen and DCS treatment cause persistence by inhibiting cell division through inhibiting peptidoglycan synthesis, while IFNγ treatment causes persistence by inducing the enzyme indoleamine-2,3-dioxygenase in the host cell which serves to deplete tryptophan levels in the cell, thus starving *Chlamydia* of this essential amino acid (37). Comparing the live cell imaging data from these different persistence inducers revealed that the IFNγ treated *Chlamydia* never expressed Clover from any promoters tested early or late. This dramatic difference in gene regulation suggests different mechanisms are likely involved and that persistence is not a phenotype associated with a specific gene expression profile.

Overall, our data support a model that RB to EB development follows a cell autonomous preprogrammed cycle that requires cell division. Our initial mathematical model assumed an inhibitory signal that at high concentrations inhibited RBs from differentiating into EBs. The concentration of this signal was depleted by metabolic utilization and RB to EB differentiation occurred. We have now updated this model to reflect our current data supporting an intrinsic signal linked to cell growth and division. This model proposes an internal signal in the nascent RB that at high concentrations inhibits RBs from differentiating into EBs, and that the signal concentration is depleted through dilution via 3-5 cell divisions and not metabolic utilization. After the inhibitory signal is reduced below a threshold, RBs are capable of transitioning to EBs (Fig. 12). Of the current proposed models in the literature, (nutrient limitation (41), inclusion membrane limitation (42), and RB size (14)) only the model based on RB size is consistent with our data. The RB size model described by Dr. Tan and colleagues at the University of California Irvine proposes that RB growth rate is lower than the division rate leading to a size reduction (depletion of signal) of the RBs after each division (14). After several rounds of division, a size threshold is reached and EB development is triggered (14). This proposed mechanism also fits our model as size would act as the inhibitory signal that is reduced through cell division. It should be noted that although we propose the dilution of an inhibitor as the intrinsic signal to control cell type switching, it is equally valid that a positive signal linked to cell division such as the development of asymmetry/polarity could act as an EB promoting signal.

**Fig. 12.**
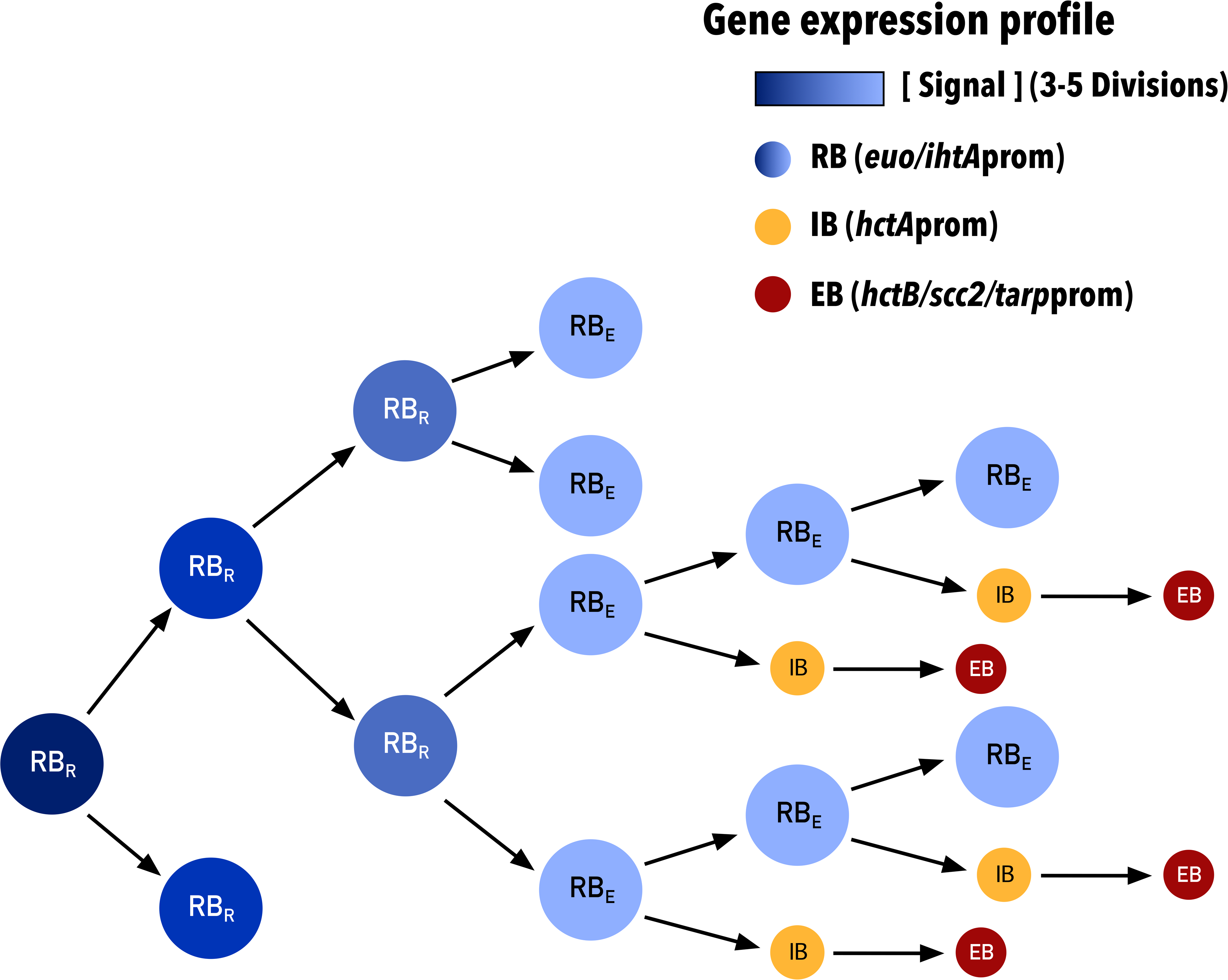
Schematic of concentration dependent RB/EB conversion model. Schematic shows diminishing signal concentration within RBs (dark to light blue) upon cell division. Depletion of the signal permits RBs to produce IBs (yellow) which then convert to EBs (red). RB_R_s divide into two subsequent RBs. RB_E_s are competent to make EBs and divide into a RB and an IB. Each cell form has predicted associated promoter expression phenotypes. RB (RB_R_ and RB_E_): *euo/ihtA*, IB: *hctA*, EB: *hctB/scc2/tarp*.

Chlamydial development can be considered to occur in two steps, an RB exponential growth step starting ~ 12 HPI (*Ctr* serovar L2) and an asynchronous EB production step starting at ~ 18 HPI (*Ctr* serovar L2) (43, 44). Although the size reduction model and our model explain some of the gene expression patterns that control cell type switching, it is clear that EB development is more complicated than these simple switch models. The output of the models fit the switch between the RB exponential growth phase and the beginning of EB development, but they do not adequately explain the continued requirement for cell division during asynchronous EB production. Our data show that Pen treatment blocks the *euo* to *hctA* gene expression switch even when added late in infection (28 HPI), well after the time of initial EB formation (~18HPI). Further evidence for a dilution independent second step is the observation that the *euo*prom to *hctA*prom switch is initially blocked by both DCS treatment and Pen treatment, yet this inhibition is eventually overcome and *hctA*prom-Clover is expressed after a 9 hour delay. Unlike Pen treatment where RBs continue to increase in size, DCS impacts cell growth resulting in smaller RBs thus limiting the effect of dilution. These observations support a second developmental regulatory step that is independent of inhibitor dilution suggesting cell division itself is an important step in committing to the EB cell type.

Our interpretation of these data is that EB formation is multifactorial and requires multiple steps to form a final infectious EB. The first step is the loss of the inhibitory signal in the RB through multiple rounds of division where RBs (RB_R_) divide 3-5 times by binary fission, eventually becoming competent to transition to an EB (RB_E_). This is followed by a second step that is dependent on asymmetric cell division creating two cells with different expression profiles. One daughter cell becomes an RB_E_ (*euo*prom positive) and the second daughter cell becomes committed to EB formation (IB, *hctA*prom positive) (Fig. 12). The committed IB cell (*hctA*prom positive) does not divide and matures into the infectious EB (*hctB*prom, *scc2*prom and *tarp*prom positive). Further divisions of the RB_E_ cell produces one RB_E_ and one IB leading to a linear increase in EBs.

Additional support for asymmetric EB production is the observation that *hctA*prom signal (IB production) follows a near perfect linear trajectory and is not logarithmic during the EB production phase (24 HPI - 40 HPI, Fig 6C and S3). This observation was also true for the *hctB*prom signal, although linear expression lagged about 5 hrs as compared to the *hctA*prom signal, again suggesting a maturation delay in EB formation. In contrast the *euo*prom signal (RB growth) transitions from log to linear to limited continued growth (Fig 6C and S3). These observations suggest that the RB_R_ cell population expands by exponential growth followed by a transition to the RB_E_ cell type. The RB_E_ then divides asymmetricity leading to EB production with no gain in RB_E_ numbers. Other studies have provided evidence for asymmetric cell division. These studies showed that the cell division machinery assembles asymmetrically potentially leading to polarized division (38, 39). Additionally, The EB itself is asymmetric demonstrating hemispherical projections that can be seen by EM (45)

Our data show that the combination of mathematical modeling and live cell gene reporter imaging is a powerful tool to tease apart the molecular details of cell type development. Continued revision and testing of our models of development will lead to an expanded understanding of cell type development in this important human pathogen.

## Materials and Methods

### Organisms and cell culture

Cos-7 and HeLa cells were obtained from (ATCC). Cos-7 cells were used for all experiments unless specified. Both Cos-7 and HeLa cells were maintained in a 5% CO2 incubator at 37°C (unless otherwise indicated) in RPMI 1640 (Cellgro) supplemented with 10% fetal plex and 10g/ml gentamicin. All *Ctr*-L2 (LGV 434) strains were grown in and harvested from Cos-7 cells. For cell division experiments chlamydial cell division was inhibited by the addition of 1 U/ml penicillin-G or 40 μg/ml D-cycloserine to the media. To starve *Chlamydia* of tryptophan HeLa cells were incubated for 24 h in media containing 2 ng/ml recombinant human IFN-γ (Invitrogen: PHC4033) prior to infection. Elementary bodies were purified by density centrifugation using 30% MD-76R 48 hours post infection (18). Purified elementary bodies were stored at −80°C in sucrose-phosphate-glutamate buffer (10 mM sodium phosphate [8 mM K 2HPO 4, 2 mM KH 2PO 4], 220 mM sucrose, 0.50 mM L-glutamic acid; pH 7.4). *E. coli* ER2925 (dam/dcm-) was utilized to produce unmethylated constructs for transformation into *Chlamydia*.

### Reporter plasmids

The backbone for all promoter-reporter constructs was p2TK2SW2 (46). Promoters were amplified from *Ctr*-L2 genomic DNA using the primers indicated in table ST1. Each promoter sequence consisted of ~100 base pairs upstream of the predicted transcription start site for the specified chlamydial genes plus the UTR and the first 30nt (10aa) of the respective ORF. Promoter sequences were inserted into p2TK2SW2 downstream of the ColE1 ORI. Fluorescent reporters (EGFP/Clover/mKate2) were ordered as gene blocks from Integrated DNA Technologies (IDT) and inserted in frame with the first 30nt of the chlamydial gene. Each ORF was followed by the incD terminator. The *bla* gene was replaced by the *aadA* gene (Spectinomycin resistance) from pBam4. The final constructs reported in this study were p2TK2*-ihtA*prom-EGFP, p2TK2-*hctA*prom-Clover, p2TK2-*hctB*prom-Clover, p2TK2-*scc2*prom-Clover, p2TK2-*euo*prom-Clover, p2TK2-*tarp*prom-Clover, p2TK2-*hctB*prom-mKate2/*euo*prom-Clover and p2TK2-*hctA*prom-mKate2/*ihtA*prom-mNeonGreen.

### Chlamydial transformation and isolation

Transformation of *Ctr*-L2 was performed as previously described (46) and selected using 500ng/ul spectinomycin. Clonal isolation was achieved via successive rounds of inclusion isolation (MOI <1) using a micro-manipulator. The plasmid constructs were purified from chlamydial transformants, transformed into *E. coli* and sequenced.

### Replating and IFUs

*hctA*prom-Clover EBs were obtained from infected Cos-7 cells by scraping the host monolayer and pelleting via centrifugation for 30 min at 17200 rcfs. The EB pellets were resuspended in RPMI via sonication. For reinfection, Cos-7 cells were plated to confluency in clear polystyrene 96-well microplates. EB reinfections consisted of 2-fold dilutions. Infections were incubated for 30 min at 37°C. Infection medium was then washed 3X with HBSS and replaced with complete RPMI 1640 + cycloheximide. Spectinomycin was added to superinfection experiments to prevent wt *Ctr*-L2 growth. Infected plates were incubated for 29 hours. Cells were fixed with methanol and stained with DAPI. The DAPI stain was used for automated microscope focus and visualization of host cell nuclei, GFP-Clover for visualization of EBs and inclusion counts. Inclusions were imaged using a Nikon Eclipse TE300 inverted microscope utilizing a scopeLED lamp at 470nm and 390nm, and BrightLine bandpass emissions filters at 514/30nm and 434/17nm. Image acquisition was performed using an Andor Zyla sCMOS in conjugation with μManager software. Images were analyzed using ImageJ software (47) and custom scripts.

### Genome number quantification

Chlamydial genomic DNA was isolated from infected host cells during active infections using an Invitrogen Purelink genomic DNA mini kit. An ABI-7900HT RT-PCR system was utilized for quantification of genomic copies. A DyNAmo Flash SYBR Green qPCR kit and hctA-specific primer was used for detection.

### Fluorescent microscopy

Cos-7 monolayers were infected with *Ctr*-L2-prom EBs and incubated for 30 min at 37°C. Infection medium was then washed 3X with HBSS and replaced with complete RPMI 1640 + cycloheximide. Live infections were grown in an OKOtouch CO2/heated stage incubator. Infections were imaged using a Nikon Eclipse TE300 inverted microscope. A ScopeLED lamp at 470nm and 595nm, and BrightLine Bandpass filters at 514/30nm and 590/20nm were used for excitation and emission. DIC was used for focus. Image acquisition was performed using an Andor Zyla sCMOS camera in conjugation with μManager software (48). Images were taken at 30 min intervals from 10 to 48 hours after *Ctr-*L2-prom infection, unless otherwise stated. Live-cell infections were performed in 24 or 96 well glass-bottom plates allowing treatments to vary between wells. Multiple fields were imaged for each treatment. Fluorescent intensities for individual inclusions were monitored over time using the Trackmate plug-in in ImageJ (22). Inclusion fluorescent intensities were then analyzed, and graphed using pandas, matplotlib, and seaborn in custom Python notebooks.

For confocal microscopy, samples were fixed with paraformaldehyde, washed with phosphate-buffered saline, and mounted with MOWIOL. Confocal images were acquired using a Nikon spinning disk confocal system with a 60x oil-immersion objective, equipped with an Andor Ixon EMCCD camera, under the control of the Nikon elements software. Images were processed using the image analysis software ImageJ (http://rsb.info.nih.gov/ij/). Representative confocal micrographs displayed in the figures are maximal intensity projections of the 3D data sets, unless otherwise noted.

## Statistical analysis

Each set of experiments was repeated a minimum of three times.

## Data Availability

All data, bacterial strains and methodologies are available upon request.

## Acknowledgments

We thank Dr. Dan Rockey at Oregon State University for supplying the isogenic *Ctr* serovar J and Js strains. This work was supported by NIH grant R01AI130072, R21AI135691 and R21AI113617. Additional support was provided by start up funds from the University of Idaho and the Center for Modeling Complex Interactions through their NIH grant P20GM104420.

## Supplemental figures

**S2. Inhibition of cell division inhibits *tarpprom* expression.** Host cells were infected with purified *Ctr-L2-tarp*prom-Clover EBs followed by treatment penicillin at 14HPI (arrow). **A.** The average of *tarp*prom-Clover (EB) expression intensities from >50 individual inclusions monitored via automated live-cell fluorescence microscopy in the absence (UNT) and presence of penicillin (Pen). Experiment was repeated four times. Cloud represents SEM.

**S3. EB production follows a linear trajectory**. Host cells were infected with purified *Ctr-*L2*-*prom EBs. **A-B.** The average fluorescent intensities from >50 individual inclusions over time expressing *euo*prom-Clover (RB) and *hctA*prom-Clover (EB) plotted on both a logarithmic plot and linear plot. **C-D.** The average fluorescent intensities from >50 individual inclusions over time expressing *euo*prom-Clover (RB) and *hctB*prom-Clover (EB) plotted on both a logarithmic plot and linear plot. Red lines are drawn to highlight the linear region of expression for *hctA*prom (A-B) and *hctB*prom (C-D) (straight line region on linear plot and curved line region on log plot). Cloud represents SEM.

**M1. Live cell time lapse movie of inclusion development and *ΛcŕA*prom-Clover expression.** Host cells were infected with purified *Ctr-*L2*-hctA*prom-Clover EBs. Automated live-cell DIC and fluorescence microscopy was used to capture images every thirty minutes from 10-48 HPI.

